# Characterizing putative glycosyltransferases within the flagella glycosylation island (FGI) of *Aeromonas hydrophila* ATCC 7966^T^

**DOI:** 10.1101/2024.06.23.600306

**Authors:** Kelly M. Fulton, Elena Mendoza-Barberà, Juan M. Tomás, Susan M. Twine, Jeffrey C. Smith, Susana Merino

## Abstract

Motility is an important virulence factor for many pathogenic bacteria, enabling locomotion towards favourable conditions and away from hostile environments. Flagellar-mediated motility is driven by one or more flagellar filaments that extend outside of the cell and rapidly rotate to generate movement. These filaments are assembled through the polymerization of thousands of copies of structural flagellin proteins. It has been shown that flagellin glycosylation is often a prerequisite for proper flagella structure and function. *Aeromonas hydrophila* ATCC 7966^T^, a clinical and environmental pathogen, elaborates a single polar flagellum. The polar flagellin structural proteins FlaA and FlaB are glycosylated with a heterologous collection of complex penta- and hexa-saccharide chains. This study characterized the involvement of four genes with homology to known glycosyltransferases located within the *A. hydrophila* ATCC 7966^T^ flagellar glycosylation island (FGI) in the biosynthesis of the complex polysaccharide glycans modifying the polar flagellins. Deletion of genes *AHA_4167*, *AHA_4169*, *AHA_4170*, and *AHA_4171* were observed to have truncated glycans with sequentially shorter chain length, and all of these mutant strains had reduced motility compared to wild type bacteria.

## 1. INTRODUCTION

Bacteria produce an array of elaborate glycoconjugate structures, such as lipopolysaccharide (LPS), capsular polysaccharide (CPS), extracellular polysaccharide (EPS), cell wall glycopolymers (CWG), and a variety of glycoproteins. Many known bacterial glycoproteins, including S-layer, pili, fimbriae, and flagella, are surface associated and interact with the external environment. While in eukaryotic organisms there are ten common monosaccharides that can be assembled into glycans, bacteria collectively produce a much wider range of unique and less well characterized carbohydrate structures that are often species or strain specific. Bacterial glycan chains can vary in length from mono-to oligosaccharide, have a linear or branched structure, be attached to the proteins by either *N-* (through asparagine) or *O-* (through serine, threonine, and sometimes tyrosine) linkage types, and involve complex biosynthetic pathways. The assembly can proceed via an oligotransferase (OTase)-dependent or OTase-independent mechanism [1,2]. The OTase-dependent mechanism involves the *en bloc* transfer of a complete glycan chain from a lipid carrier onto a target protein in the periplasm. By contrast, the OTase-independent mechanism involves the sequential addition of individual carbohydrate moieties directly onto the target protein by a collection of specific glycosyltransferase (GT) enzymes in the cytoplasm.

In pathogenic bacteria, protein glycosylation has been associated with a variety of key virulence factors, such as motility, host recognition, adherence, colonization, and immune evasion. Flagellar-mediated motility of bacterial pathogens, for example, often depends upon glycosylation of the structural flagellin proteins. In the absence of flagellin glycosylation, many species are incapable of assembling the flagellar filament and motility is consequently reduced or eliminated. This has been demonstrated in both Gram-positive (i.e. *Clostridium difficile* [3], *Clostridium botulinum* [4], *Listeria monocytogenes* [5]) and Gram-negative (i.e. *Campylobacter jejuni* [6], *Burkholderia thailandensis* [7]*, Burkholderia pseudomallei* [7]*, Aeromonas piscicola* [8] bacteria. Furthermore, disruption to or alteration of many flagellin glycans has been linked to changes in the affinity with which a bacterium can recognize and bind to host cells. For example, in *C. jejuni*, the acetamidino form of pseudaminic acid (PseAm) was required for virulence in the ferret model of pathogenesis [9] and the loss of the acetamidino form of legionaminic acid (AmLeg) reduced colonization of chickens [10].

Mesophilic *Aeromonas* species are Gram-negative pathogens that are ubiquitous in aquatic environments. They are capable of infecting a range of fish and aquatic animals, with motile aeromonad septicemia being the most common presentation. They are also zoonotic, commonly causing infections of the human gastrointestinal tract [11] following consumption of contaminated food or water and soft tissue infections through open wounds [12]. Exacerbating the clinical concern is an observed increase in multidrug resistant (MDR) and extremely drug resistant (XDR) strains. These bacteria possess inducible lateral flagella but are primarily motile in liquid media via a single constitutively expressed polar flagellum. The polar flagellar filament is an oligomer of two structural proteins, FlaA and FlaB. These proteins are encoded by the *flaA* and *flaB* genes, which are located in region 2 of the flagellum biosynthesis gene clusters [13,14]. Also located in region 2 is a gene with homology to motility accessory factor 1 (*maf-1*), which is putatively a glycosyltransferase that adds a nonulosonic linking sugar, such as pseudaminic acid (Pse), to flagellin proteins [15]. Bioinformatic analysis has also revealed the existence of flagella glycosylation islands (FGIs) flanked by genes homologous to *pse* biosynthesis genes in 210 of 265 *Aeromonas* genomes evaluated [16]. These FGIs are often, though not always, downstream of the flagellum biosynthesis region 2. They are highly polymorphic across species but have been categorized into three groups. Group I and III FGIs are relatively small and are associated with strains that modify flagellin with single monosaccharides. By contrast, Group II FGIs are larger and encode a collection of putative glycosyl-, phospho-, methyl-, and acetyl-transferases as well as proteins involved in fatty acid synthesis and a variable number of open reading frames (ORFs) with no known homology. All strains with Group II FGIs, except *Aeromonas caviae* strain Aer 268, contain a gene with homology to the *A. hydrophila* strain AH-3 *fgi-1* glycosyltransferase, which has been shown in that strain to add hexose (Hex), the second carbohydrate of the glycan, to the Pse linking sugar [16]. A gene with homology to motility accessory factor 2 (*maf-2*) is also present adjacent to most Group II FGIs.

The genome of *Aeromonas hydrophila* ATCC 7966^T^, which was published in 2006 [17], contained a 27 kb Group II FGI which represents 31 ORFs, and therefore suggested that the FlaA and FlaB proteins would be modified with a complex glycan chain. This was confirmed by mass spectrometry analysis that revealed these flagellins were modified with diverse penta- and hexa-saccharide glycans [18] that had variable secondary modification with phosphate and methyl groups. The glycan had a novel 422 Da Pse derivative (Pse_d_) linking sugar, followed in sequence by two Hex, a putative *N*-acetylglucosamine (GlcNAC) with variable secondary modification, a putative deoxy-GlcNAc (dGlcNAc) or methylated-dGlcNAC, and at a subset of sites, an additional GlcNAc.

This study aimed to improve understanding of *A. hydrophila* ATCC 7966^T^ polar flagellin glycan assembly by evaluating the role of four putative glycosyltransferase genes encoded in the FGI. The genes were deleted and polar flagellin proteins from each of the corresponding mutants were purified. Flagellin glycosylation was analyzed using mass spectrometry, with data suggesting that each gene product had a specific role in glycan assembly beyond the Pse_d_ linking sugar. The flagellin glycans observed in the ATCCΔAHA_4171 (denoted *fgi-1* in *A. piscicola* AH-3), ATCCΔAHA_4170, ATCCΔAHA_4169, and ATCCΔAHA_4167 mutant strains were truncated predominantly to mono-, di-, tri-, and tetra-saccharides, respectively, as compared to the penta- and hexa-saccharides observed in the wild type strain. Furthermore, less extensive glycosylation correlated with poor flagellum assembly and reduced motility.

## 2. METHODS

### 2.1 Bacterial strains, plasmids, and growth conditions

Bacterial strains and plasmids used in this study are listed in **Table 1**. *A. hydrophila* ATCC 7966^T^ and its mutants were grown in tryptic soy broth (TSB) or agar (TSA) at 30 °C. *Escherichia coli* strains were grown on Luria-Bertani (LB) Miller broth and LB Miller agar at 37 °C. When required, chloramphenicol (25 µg/mL), rifampicin (100 µg/mL) and spectinomycin (50 µg/mL) were added to media.

**Table 1.** Bacterial strains and plasmids used in this study.

| Strain or plasmid | Genotype and/or phenotype | Reference |
| --- | --- | --- |
| **Strains** |  |  |
| ***A. hydrophila*** |  |  |
| ATCC7966<sup>T</sup> | *A. hydrophila* wild type | [17] |
| ATCC7966-Rif | ATCC7966<sup>T</sup>, spontaneous Rif<sup>r</sup> | [19] |
| ATCCΔAHA4171 | ATCC7966-Rif; ΔAHA_4171 | [16] |
| ATCCΔAHA4170 | ATCC7966-Rif; ΔAHA_4170 | This work |
| ATCCΔAHA4169 | ATCC7966-Rif; ΔAHA_4169 | This work |
| ATCCΔAHA4167 | ATCC7966-Rif; ΔAHA_4167 | This work |
| ***E. coli*** |  |  |
| HB101 | *F- mcrB mrr hsdS20(rB- mB-) recA13 leuB6 ara-14 proA2lacY1 galK2 xyl-5 mtl-1 rpsL20(SmR) glnV44 λ* | [20] |
| MC1061λpir | *thi thr1 leu6 proA2 his4 argE2 lacY1 galK2 ara14 xyl5 supE44 λ pir* | [21] |
| **Plasmids** |  |  |
| pRK2073 | Helper plasmid, Sp<sup>r</sup> | [21] |
| pDM4 | Suicide plasmid, *pir* dependent with *sacAB* genes, oriR6K, Cm<sup>r</sup>. | [22] |
| pDM-AHA4171 | pDM4ΔAHA_4171 of ATCC7966<sup>T</sup>, Cm<sup>r</sup>. | [16] |
| pDM-AHA4170 | pDM4ΔAHA_4170 of ATCC7966<sup>T</sup>, Cm<sup>r</sup>. | This work |
| pDM-AHA4169 | pDM4ΔAHA_4169 of ATCC7966<sup>T</sup>, Cm<sup>r</sup>. | This work |
| pDM-AHA4167 | pDM4ΔAHA_4167 of ATCC7966<sup>T</sup>, Cm<sup>r</sup>. | This work |
| pBAD33 | pBAD33 arabinose-induced expression vector with Cmr | [23] |
| pBAD33-AHA4171 | pBAD33 with AHA_4171 of ATCC7966<sup>T</sup>, Cmr | [16] |
| pBAD33-Fgi-1 | pBAD33 with fgi-1 of AH-3, Cmr | [16] |

Rif<sup>r</sup>, rifampicin resistant; Cm<sup>r</sup>, chloramphenicol resistant; Sp<sup>r</sup>, spectinomycin resistant.

### 2.2 Construction of ATCC 7966^T^ glycosyltransferase mutants

Single defined in-frame deletion of ATCC 7966^T^ putative glycosyltransferases genes *AHA_4171*, *AHA_4170*, *AHA_4169* and *AHA_4167* were obtained by allelic exchange [22] using primers listed in **Table 2**. Construction of pDM-AHA4171 plasmid and generation of ATCCΔAHA4171 mutant was previously described [16]. Briefly, deletions were performed by amplification of DNA regions upstream (fragment AB) and downstream (fragment CD) of each glycosyltransferase of *A. hydrophila* ATCC 7966^T^ in two sets of asymmetric polymerase chain reactions (PCRs). Primer pairs A-4170 and B-4170, and C-4170 and D-4170, amplified DNA fragments of 733 bp (AB-4170) and 696 bp (CD-4170) upstream and downstream of AHA_4170, respectively. Primer pairs A-4169 and B-4169, and C-4169 and D-4169, amplified DNA fragments of 749 bp (AB-4169) and 637 bp (CD-4169) upstream and downstream of AHA_4169, respectively. Primer pairs A-4167 and B-4167, and C-4167 and D-4167, amplified DNA fragments of 789 bp (AB-4167) and 663 bp (CD-4167) upstream and downstream of AHA_4167, respectively. DNA fragments AB / CD-4170, AB / CD-4169, and AB / CD-4167 were annealed at their overlapping regions and amplified as a single fragment using primers A-4170 and D-4170, A-4169 and D-4169, and A-4167 and D-4167, respectively. The AD fusion products were purified, *Bgl*II or *Bam*HI digested, ligated into *BglII*-digested and phosphatase-treated pDM4 vector [22] and electroporated into *E. coli* MC1061 (*λpir*). Recombinant plasmids pDM-AHA4170, pDM-AHA4169 and pDM-AHA4167 were selected on chloramphenicol plates at 30 °C and introduced into *A. hydrophila* ATCC 7966^T^ rifampicin-resistant (ATCC 7966-Rif) by triparental mating using the *E. coli* MC1061 *(λpir*) containing the insertion constructs and the mobilizing strain HB101/pRK2073. Transconjugants were selected on chloramphenicol and rifampicin plates. After sucrose treatment, rifampicin-resistant (Rif^r^) and chloramphenicol sensitive (Cm^S^) transformants were chosen and single defined in-frame deletion were confirmed by PCR.

**Table 2.**
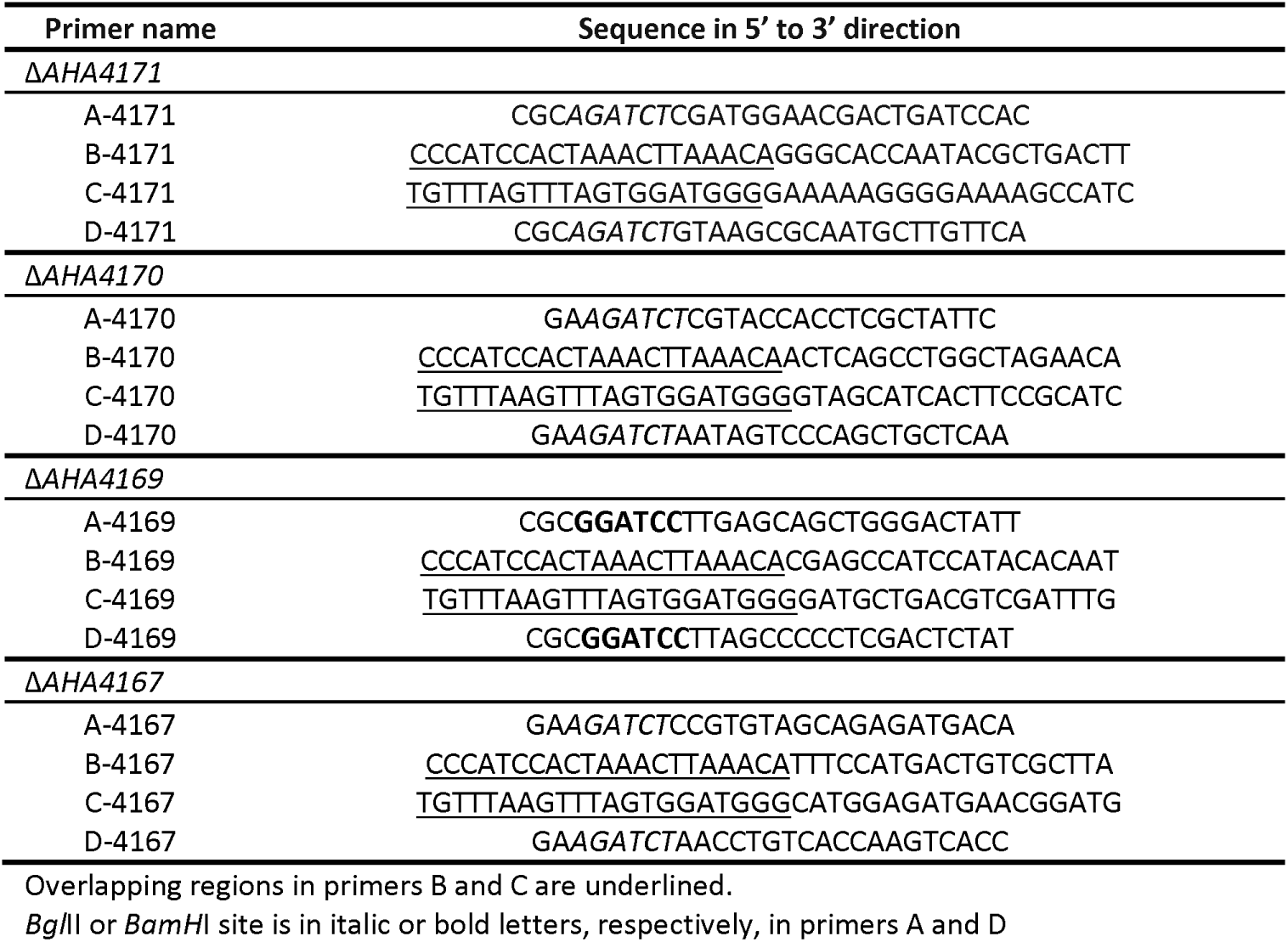
Primers used for the construction of the in-frame defined mutants.

### 2.3 General DNA techniques and nucleotide sequencing

DNA manipulations were carried out according to standard procedures [24] and as recommended by the supplier of products used in each procedure. Restriction endonucleases, T4 DNA ligase and alkaline phosphatase were obtained from Invitrogen. PCRs were performed in a Gene Amplifier PCR System 2400 Perkin Elmer Thermal Cycler, using AccuPrime™ Taq DNA polymerase (Invitrogen). Standard PCR analysis was performed using Dream Taq DNA polymerase (Thermo Scientific^TM^). Plasmid DNA for sequencing was isolated by GeneJet Plasmid Miniprep Kit (Thermo Scientific^TM^) as recommended by the suppliers. DNA sequencing was performed with the BigDye Terminator v3.1 cycle sequencing kit (Applied Biosystem). Custom-designed primers were purchased from Sigma-Aldrich.

### 2.4 Identification and characterization of glycosyltransferases

Genome sequences were retrieved from the National Center for Biotechnology (NCBI) database and identities of deduced amino acid sequences were inspected in the Gene Bank database using the BLASTP network service at NCBI. Protein domains were determined with the NCBI Conserved Domain Database (CDD) [25], and the SMART sequence analysis [26]. The presence of signal peptides was explored with the SignalP 4.1 Server [27] and of transmembrane helices using the TMHMM Server v2.0 [28]. Secondary structure prediction was executed with SOPMA [29] and tertiary structure was modeled with the Intensive method from the Phyre2 Protein Fold Recognition Server [30].

### 2.5 Flagellin Purification

Wild type and mutant strains of *A. hydrophila* ATCC 7966^T^ were grown in tryptic soy broth (TSB) at 30 °C overnight. Cells collected by centrifugation at 5000 x *g* and resuspended in 100 mM Tris (pH 7.8). Polar flagella were stripped from bacterial cells using mechanical shearing and then purified, as described previously [8,18].

### 2.6 Motility assays

To assess the swimming motility of wild type and mutant strains of *A. hydrophila* ATCC 7966^T^, fresh bacterial colonies were transferred with a sterile toothpick onto the center of a soft agar plate (1% tryptone, 0.5% NaCl, 0.25% agar). Plates were incubated face up at 25 °C for 18-24 h, and motility was assessed by examining the migration of bacteria through the agar from the center towards the periphery of the plate. Moreover, swimming motility was assessed by light microscopy observations in liquid media.

### 2.7 SDS-PAGE

Polar flagellins purified from bacterial growths of each mutant strain (2 µg) were separated by 12% SDS-PAGE as described previously [31] and stained with Coomassie Blue total protein stain (BioRad), according to the manufacturer’s instructions.

### 2.8 Mass spectrometry analysis

Purified polar flagellin proteins were digested with trypsin as described previously [18]. Nano liquid chromatography tandem mass spectrometry (nLC-MS/MS) was performed using an Ultimate 3000 LC (Dionex) coupled to an Orbitrap Exploris 480 mass spectrometer (Thermo Fisher). The nLC separation peptides was performed exactly as described previously [18]. MS^1^ spectra were acquired between m/z 350-1600 at a resolution of 60,000. Using data-dependent acquisition (DDA), MS^2^ fragmentation achieved by high voltage collision-induced dissociation (HCD) with a normalized collision energy of 30% and spectra were acquired in the orbitrap with a resolution of 60,000. Glycopeptide spectra were manually *de novo* sequenced.

## 3. RESULTS

### 3.1 Genomic identification of putative *A. hydrophila* ATCC 7966^T^ polar flagella glycosyltransferase enzymes within the FGI

The polar flagella glycosylation island (FGI) of *A. hydrophila* ATCC 7966^T^ is complex and belongs to group II, like the FGI of *A. piscicola* AH-3 [16]. Unlike the FGI of *A. piscicola* AH-3, whose FGI region is immediately downstream of polar flagellum region 2, the FGI of *A. hydrophila* ATCC 7966^T^ is 2,731.874 kb downstream of the polar flagellum region 2 and transcribed in the opposite direction (**Fig. 1a**). In *A. hydrophila* ATC 7966^T^, the pseudaminic acid biosynthesis genes are separated by a fragment of 27 kb, with low % G+C content (47.76%), that encodes 25 ORFs (**Fig. 1b**). Analysis of the encoded proteins using the BLASTP network service of NCBI showed that *AHA_4172* to *AHA_4176* ORFs encode proteins involved in fatty acid synthesis and *AHA_4152* to *AHA_4171* ORFs encode glycosyl-, methyl-, acetyl-, and phospho-transferases [18]. Only four of these genes encode putative glycosyltransferases: *AHA_4171*, *AHA_4170*, *AHA_4169*, and *AHA_4167*.

**Figure 1:**
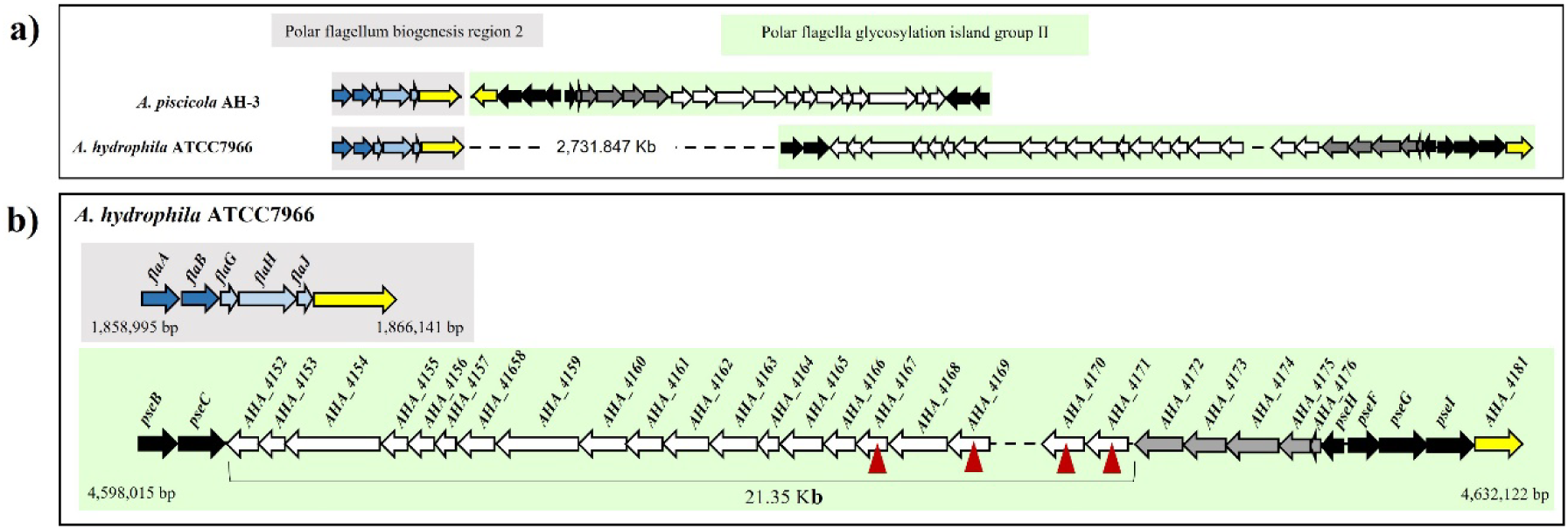
Genetic organization of polar flagellum biogenesis region 2 and polar flagella glycosylation island (FGI) group II. a) Chromosomal location and gene transcription of the flagellum biogenesis region 2 (grey box) and polar FGI group II (green box) *A. piscicola* AH-3 and *A. hydrophila* ATCC 7966^T^ . b) The grey box shows the polar flagellum biogenesis region 2 of *A. hydrophila* ATCC 7966^T^ which contained genes *flaA* and *flaB* (dark blue arrows), encoding for the structural flagellin proteins FlaA and FlaB. The green box shows the flagella glycosylation island (FGI) of *A. hydrophila* ATCC 7966^T^, with genes homologous to pseudaminic acid biosynthesis (black arrows) at either end as well as FGI genes (white arrows). Within the FGI there was a 21.35 kb region with 20 ORFs. This region contained four ORFs with high homology to glycosyltransferase enzymes: *AHA_4167, AHA_4169, AHA_4170,* and *AHA_4171* that were single in-frame deleted (red triangle) for this study. Motility accessory factor *maf-1* and *maf-2* (yellow arrows), and lipid biosynthesis gene (grey arrows) are also indicated.

The *AHA_4171* and *AHA_4167* encoded protein sequences that have 318 and 277 amino acid residues, a predicted molecular weight of 36.89 and 32.71 kDa, and a predicted pI of 5.94 and 5,02, respectively. According to CDD and SMART, both proteins showed a glycosyltransferase family 2 (GT-2) conserved domain in the first 175 amino acids, containing in the first 100 amino acids, 5 active sites characteristics of glycosyltransferase family A (GT-A). The *AHA_4170* encoded a protein sequence that has 366 amino acid residues, a predicted molecular weight of 41.73 kDa, and a predicted pI of 6.34. According to CDD and SMART, this protein showed a glycosyltransferase family 8 (GT-8) conserved domain in the first 237 amino acids, with interval 100-250 containing the 3 metal binding sites characteristic of this domain. The *AHA_4169* encoded protein sequence has 378 amino acid residues, a predicted molecular weight of 43.24 kDa, and a predicted pI of 6.04. This protein, according to CDD and SMART, showed a glycosyltransferase family 1 (GT-1) conserved domain in the interval amino acids 192-309, with glycosyltransferase family B topology (GT-B) (**Fig. 2**).

**Figure 2:**
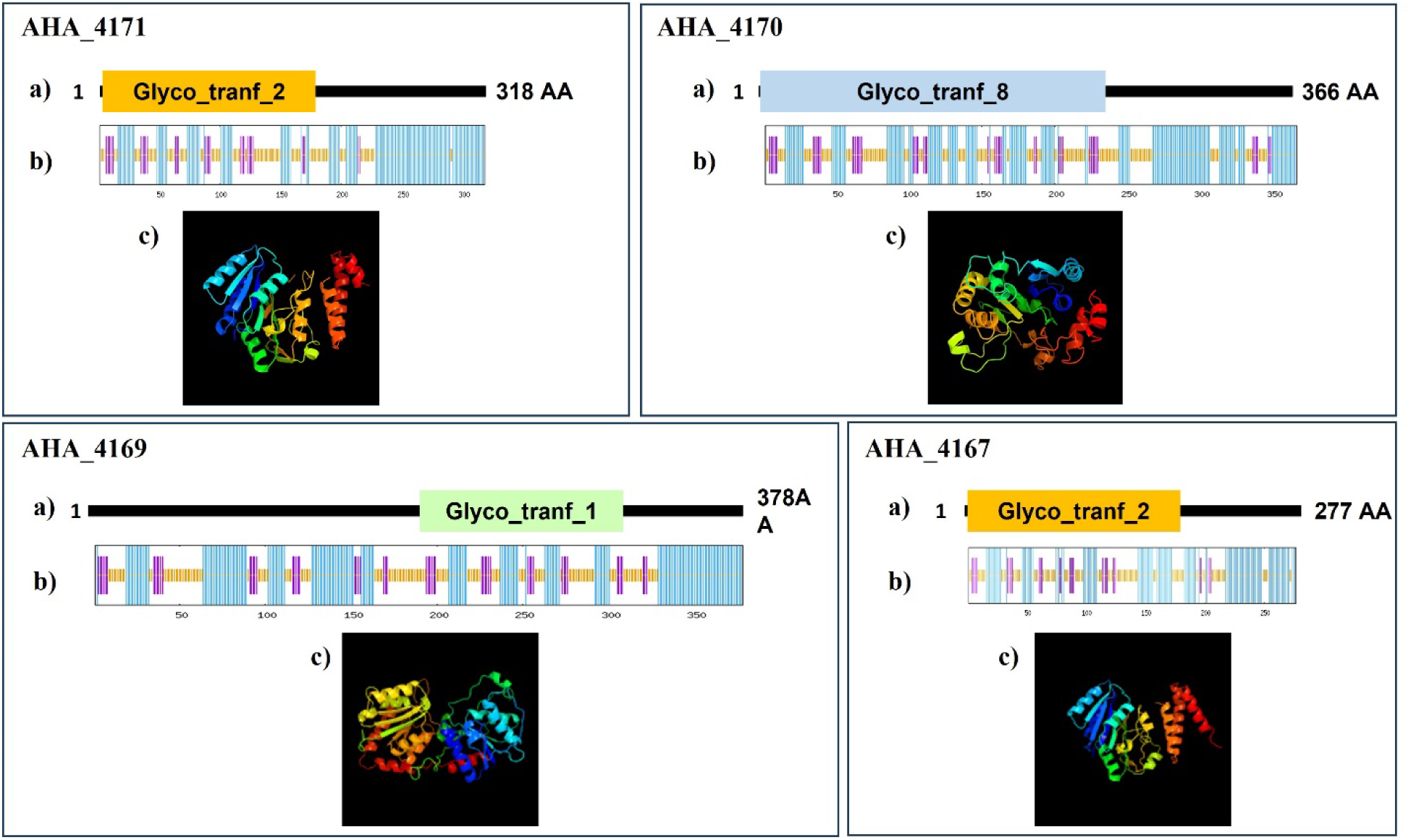
Characteristics of *A. hydrophila* ATCC 7966. ^T^ **FGI glycosyltransferases** . a) Protein domains determined with the NCBI Conserved Domain Database (CDD) and the SMART sequence analysis. b) Secondary structure predicted with SOPMA. Helix (Blue lines), coil (orange lines) and sheet (purple lines). c) Tertiary structure modeled with Phyre2 Protein Fold Recognition Server.

None of the four proteins encoded by genes with homology to glycosyltransferase enzymes had a signal peptide or transmembrane domains according to SignalP-HMM v4.1 and TMHMM Server v2.0, suggesting that they are active in the bacterial cytoplasm. All of them had orthologues in the FGI cluster of several other species and strains belonging to the FGI IIB group, such as *A. hydrophila* strains TPS30 and NCTC8049 and *Aeromonas dhakensis* strains KN-Mc-6U21 and SSU. Furthermore, the *AHA_4171* encoded a conserved glycosyltransferase downstream of *luxC*, that also showed 70% similarity and 55% identity to the Fgi-1 glycosyltransferase of *A. piscicola* AH-3 (OCA61121.1) belonging to the FGI subgroup IIA [16].

Secondary structure of these four glycosyltransferases analyzed with SOPMA predicted that they were predominantly alpha helices (51.89 to 46.56%) and random coils (40.07 to 36.79%), with a minor part of the structure predicted to be extended strands (14.81 to 11.32%). Finally, the tertiary structure of these proteins was studied using homology modelling with Phyre2 (**Fig. 2**). This approach modelled most of the tertiary structure based on the conformation of available transferase enzymes. Phyre2 used the PDB transferase c5heaA as a template to model 83% (264 residues) of AHA_4171 protein (ABK38423.1) and the 98% (271 residues) of AHA_4167 protein (ABK37536.1) with 100 and 99.97% confidence and 22 and 15% identity, respectively. To model the last residues of AHA_4171 protein, different templates with >99% confidence and sequence identity of 21 to 15% were used. The PDB transferase c6u4bA was used as template to model the 72% (262 residues) of AHA_4170 protein (ABK36197.1) with 100% confidence and 23% identity. The PDB c2qzsA transferase was used to model the 96% (362 residues) of AHA_4169 protein (ABK3640.1) with 100% confidence and 15% identity (**Fig. 2**).

### 3.2 Functional characterization of putative FGI glycosyltransferases of *A. hydrophila* ATCC 7966^T^

Based on the known amino acid sequences, the predicted molecular weights of *A. hydrophila* ATCC 7966^T^ flagellin proteins FlaA and FlaB were both approximately 32 kDa. Fully glycosylated polar flagellin proteins purified from wild type bacteria migrated to a molecular mass of 45 kDa by SDS-PAGE, as previously described. We had demonstrated that chromosomal region containing AHA_4152 to AHA_4171 of *A. hydrophila* ATCC 7966^T^ was involved in the post-translational modification of polar flagellins [18]. In order to elucidate the role of four putative glycosyltransferases within the FGI cluster of *A. hydrophila* ATCC 7966^T^, we constructed specific in-frame mutants in *AHA_4171* (ATCCΔAHA4171), *AHA_4170* (ATCCΔAHA4170), *AHA_4169* (ATCCΔAHA4169) and *AHA_4167* (ATCCΔAHA4167) using the suicide plasmids pDM-AHA4171, pDM-AHA4170, pDM-AHA4169 and pDM-AHA4167, respectively (**Fig. 1b**). Specific deletion of genes was confirmed by PCR and sequence analysis. The four mutants show reduced swimming motility in liquid medium, by light microscopy, and decreased radial expansion, ranging to 63 to 43%, in soft agar in relation to the wild type strain (**Fig. 3**).

**Figure 3.**
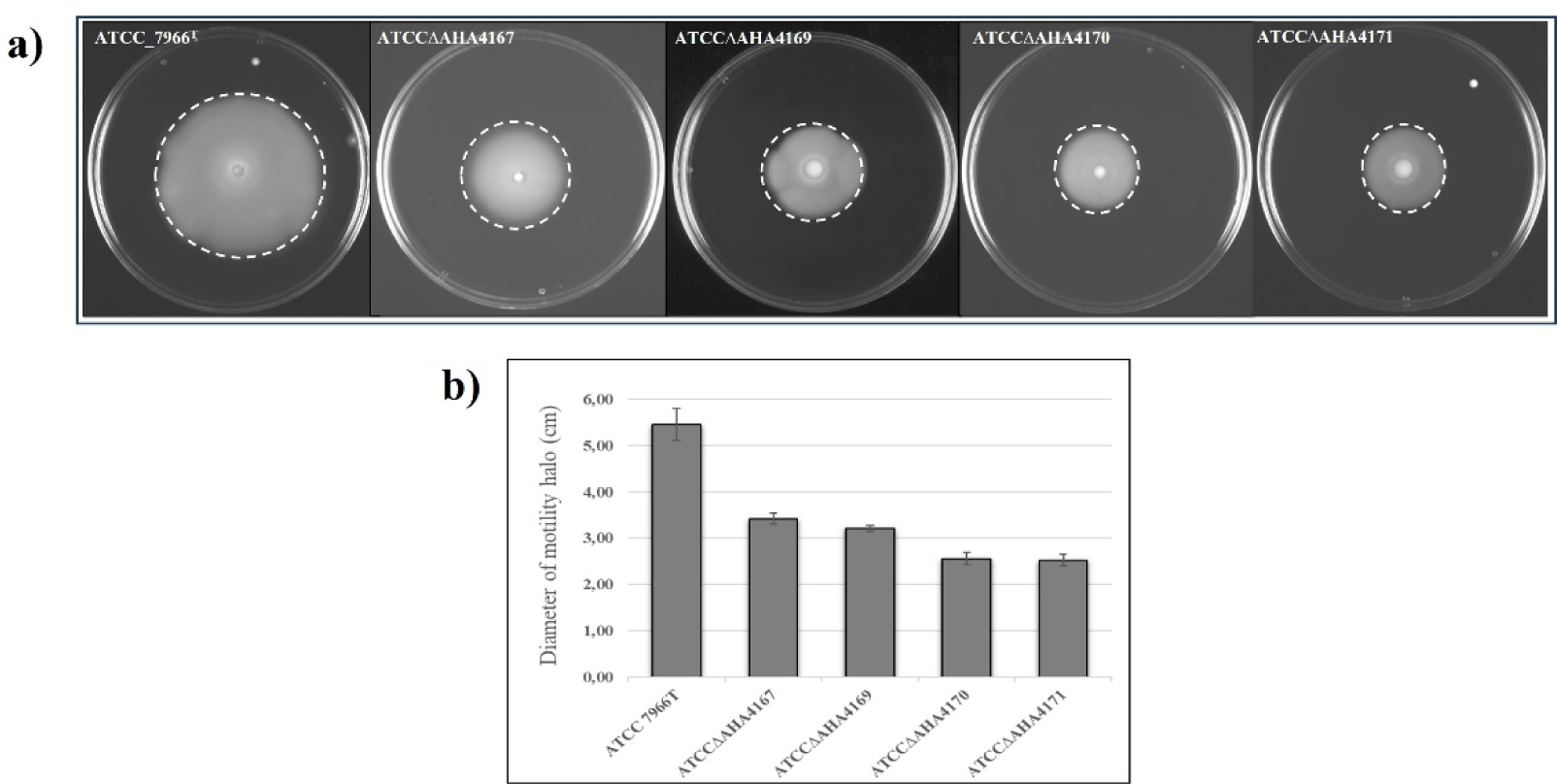
Motility of *A. hydrophila* ATCC 7966^T^ and mutants ATCCΔAHA4167, ATCCΔAHA4169, ATCCΔAHA4170, and ATCCΔAHA4171. a) Bacteria were grown for 24h at 30 °C on soft agar plates. b) The diameter of each motility halo was determined and average measurements are presented (n=6) with the standard error indicated.

Furthermore, the analysis of purified polar flagella showed that the polar flagellins of these four mutants migrated to sequentially lower molecular weights in 12% SDS-PAGE compared to the wild type, suggesting less extensive post-translational modification (**Fig. 4**).

**Figure 4:**
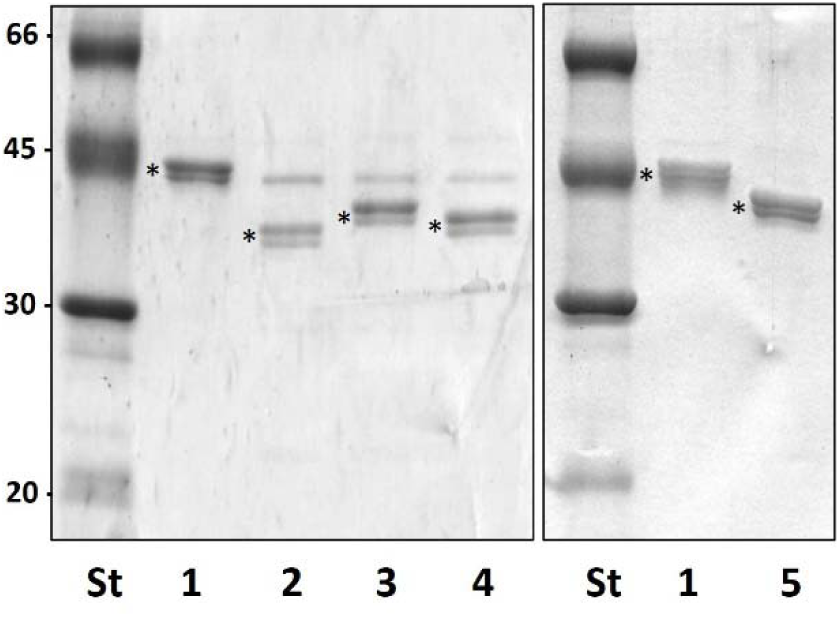
SDS-PAGE of *Aeromonas hydrophila* ATCC 7966T glycosyltransferase mutants. Purified polar flagellum of wild type ATCC 7966^T^ (lane 1), ATCCΔAHA4171 (lane 2), ATCCΔAHA4169 (lane 3), ATCCΔAHA4170 (lane 4), and ATCCΔAHA4167 (lane 5). Mutants showed a progressively greater reduction in molecular weight compared to the wild type. Polar flagellins are annotated (*) and molecular weight standards (St) are indicated (kDa).

### 3.3 Mass spectrometry analysis of putative glycosylation mutants

Peptides from tryptic digests of polar flagellin proteins purified from wild type and putative glycosyltransferase mutant strains were analyzed by nLC-MS/MS. The complex glycosylation of the polar flagellin proteins FlaA and FlaB in wild type bacteria has been previously reported [18]. The six unique FlaA and FlaB peptides were modified with collection of pentasaccharides: Pse_d_, followed by two Hex, a putative GlcNAc that was variably secondarily modified with phosphate and methyl groups, and finally a putative dGlcNAc or a MeGlcNAc terminal sugar. There was a pair of isobaric peptides, with one sequence belonging to FlaA and the other belonging to FlaB. These isobaric peptides were uniquely modified with a hexasaccharide glycan as the result of an additional HexNAc as the terminal sugar.

With respect to the non-isobaric peptides, the glycosylation of flagellin isolated from the mutant strains was significantly less complex than that of wild type bacteria, and all glycans were truncated. For flagellin purified from the ATCCΔAHA4167, ATCCΔAHA4169, ATCCΔAHA4170, and ATCCΔAHA4171 mutant strains, only a tetra, tri, di, and monosaccharide modification were observed, respectively. For simplicity, the 1233.62 Da FlaB GSATVQSVATADK sequence (T^174-186^) was selected as a representative peptide to demonstrate this glycosylation pattern. Combined MS^1^ spectra from 9.7-10.4 min permitted comparison of the glycosylation status in each mutant strain to wild type bacteria. For wild type bacteria (**Fig. 5a**), the combined MS^1^ spectrum revealed dozens of peaks representing the isobaric flagellin peptides modified with a collection of heterologous glycans up to pentasaccharide length, as previously reported [18]. The equivalent combined MS^1^ spectra for the mutant strains each contained a single peak representing the peptide modified with a truncated glycan. For the ATCCΔAHA4167 mutant strain (**Fig. 5b**), the peak at m/z 1099.48^2+^ corresponded to the FlaB peptide with a tetrasaccharide (Pse_d_-Hex-Hex-MeHexNAc) lacking the fifth carbohydrate moiety, and was the longest glycan observed. Furthermore, the fourth position of the glycan was only occupied by a MeHexNAc and there was no evidence of the highly variable secondary phosphorylation and methylation that existed in the wild type. For the ATCCΔAHA4169 mutant strain (**Fig. 5c**) the peak at m/z 990.95^2+^ represented the peptide modified with a trisaccharide (Pse_d_-Hex-Hex), lacking the fourth and fifth carbohydrate moieties, and was the longest observed glycan. For the ATCCΔAHA_4170 mutant strain (**Fig. 5d**) the peak at m/z 909.92^2+^ corresponded to the peptide modified with only a disaccharide (Pse_d_-Hex), which was the longest observed glycan. Finally, the combined MS^1^ spectrum for the ATCCΔAHA_4171 mutant strain (**Fig. 5e**) produced single peak at m/z 828.90^2+^ which corresponded to the peptide modified with a single monosaccharide (the 422 Da pseudaminic acid derivative). There was also a minor peak at less than 10% relative intensity corresponding to the peptide modified with an alternate 376 Da Pse_d_ monosaccharide. There was no evidence that the glycans modifying flagellins of the ATCCΔAHA4171, ATCCΔAHA4170, ATCCΔAHA4169, and ATCCΔAHA4167 mutant strains could exceed a mono-, di, tri, or tetra-saccharide, respectively, in contrast to the pentasaccharides modifying the flagellins of wild type bacteria.

**Figure 5:**
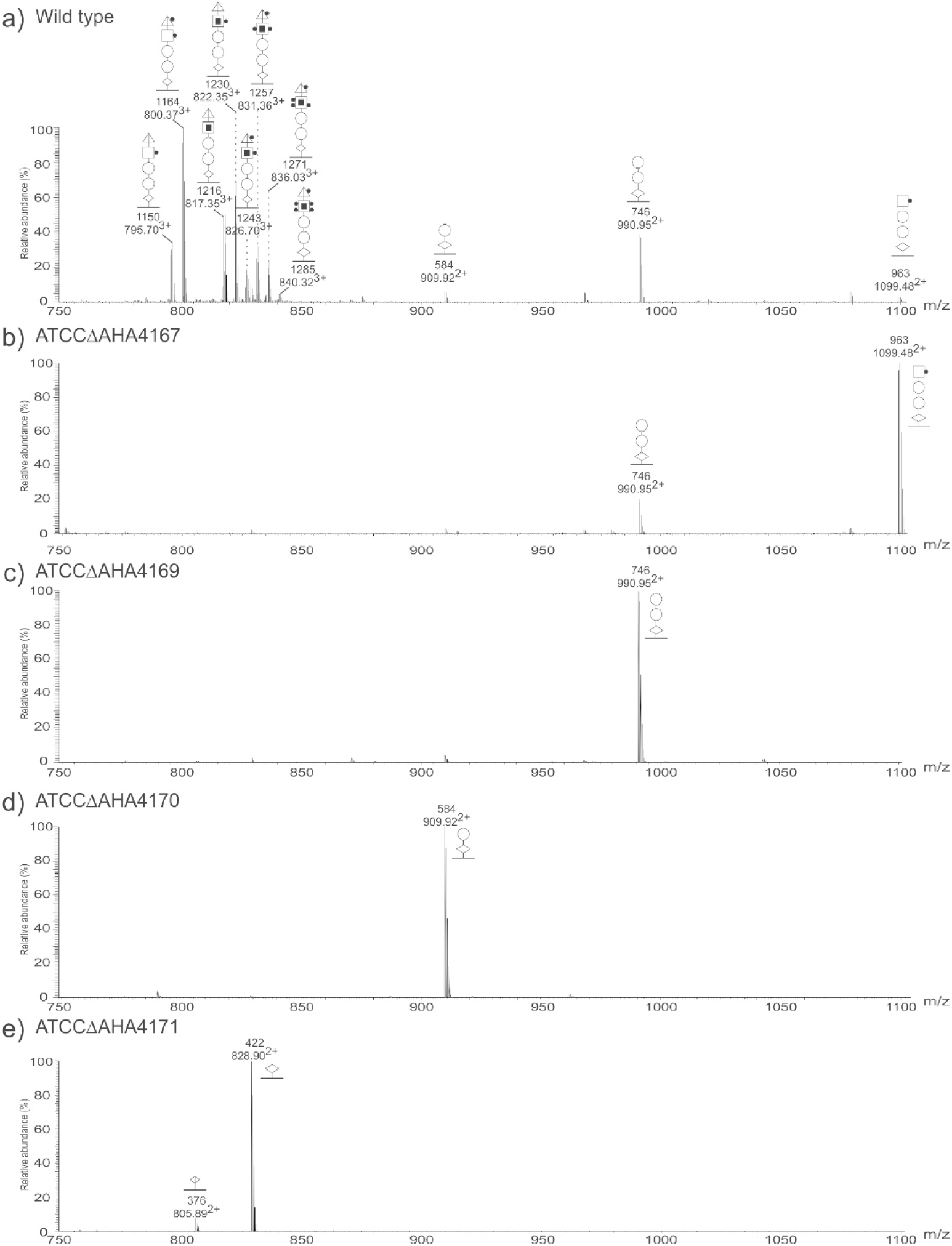
Demonstrating the impact of putative glycosyltransferase enzymes on polar flagellin glycan assembly with nLC-MS. Shown is the combined nLC-MS spectrum between 21.8-22.3 min with peaks representing the 1233.62 Da FlaB T^159-166^ peptide and associated glycosylation. The glycosylation pattern was representative of the six unique FlaA and FlaB peptides sequences. Bacteria from wild type (a) modified these peptides with a heterologous set of pentasaccharide glycans. Bacteria from mutant strains ATCCΔAHA4167 (b), ATCCΔAHA4169 (c), ATCCΔAHA4170 (d), and ATCCΔAHA4171 (e) do not extend the glycan beyond positions four, three, two, and one, respectively. In each mutant, however, the truncated glycan is also observed with an additional HexNAc residue. The additional HexNAc capping sugar was only observed in association with the isobaric peptides. Glycan masses (Da) are indicated above the glycopeptide ion m/z. 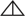: dHexNAc, 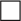: HexNAc, 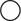: Hex, 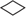: 422 Da pseudaminic acid derivative, 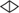: 376 Da pseudaminic acid derivative, 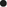:Methyl, 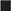: Phosphate, 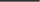: Peptide

The glycosylation pattern of isobaric FlaA/FlaB glycopeptides (T^94-108^) varied from that of the other six glycopeptides. For wild type bacteria (**Fig. 6a**), the combined MS^1^ spectrum between 7.5 and 8.5 min revealed numerous peaks representing the isobaric peptides modified with a collection of heterologous penta- and hexa-saccharide glycans, as previously reported [18]. The hexasaccharide was consistently capped with a HexNAc residue. The equivalent combined MS^1^ spectra for the mutant strains each contained a peak representing the peptide modified with a truncated glycan in the same pattern as the non-isobaric peptides. However, in each spectrum, there was a second glycopeptide species that was in all cases the truncated glycan further modified with an additional HexNAc terminal sugar, extending the chain length by one moiety. For the ATCCΔAHA4167 mutant strain (**Fig. 6b**), the combined MS^1^ spectrum revealed peaks at m/z 891.03^3+^ and m/z 958.72^3+^. The first peak corresponded to the isobaric peptides modified with a tetrasaccharide (Pse_d_-Hex-Hex-MeHexNAc) glycan lacking the fifth carbohydrate typically observed with respect to wild type. The second peak represented the peptides modified by a pentasaccharide that differed from the wild type pentasaccharide. In this case, the glycan was in fact the truncated tetrasaccharide plus an additional HexNAc capping sugar instead of the dHexNAC that was associated with wild type (i.e. Pse_d_-Hex-Hex-MeGlcNAc-HexNAc instead of Pse_d_-Hex-Hex-MeGlcNAc-dHexNAc). The extensive secondary modification of the fourth carbohydrate with methyl and phosphate groups that was documented for wild type flagellin glycosylation was not observed in this mutant strain. The combined MS^1^ spectrum for the ATCCΔAHA4169 mutant strain (**Fig. 6c**) also produced two peaks, with one at m/z 818.68^3+^ and the other at m/z 886.37^3+^. The first peak represented the peptide modified with a trisaccharide (Pse_d_-Hex-Hex), while the second peak corresponded to the peptides modified with the trisaccharide and an additional HexNAc capping sugar (Pse_d_-Hex-Hex-HexNAc). The combined MS^1^ spectrum for the ATCCΔAHA4170 mutant strain (**Fig. 6d**) similarly revealed two peaks, with one at m/z 764.66^3+^ and the other at m/z 832.35^3+^, which corresponded to the same peptides modified with a disaccharide (Pse_d_-Hex) and a disaccharide plus an additional HexNAc capping sugar (Pse_d_-Hex-HexNAc), respectively. Finally, the combined MS^1^ spectrum for the ATCCΔAHA4171 mutant strain (**Fig. 6e**) revealed two peaks with one at m/z 710.64^3+^ and the other at m/z 778.33^3+^. These peaks were shown to be the same peptides modified with a single monosaccharide (the 422 Da Pse_d_) and a disaccharide (Pse_d_-HexNAc).

**Figure 6:**
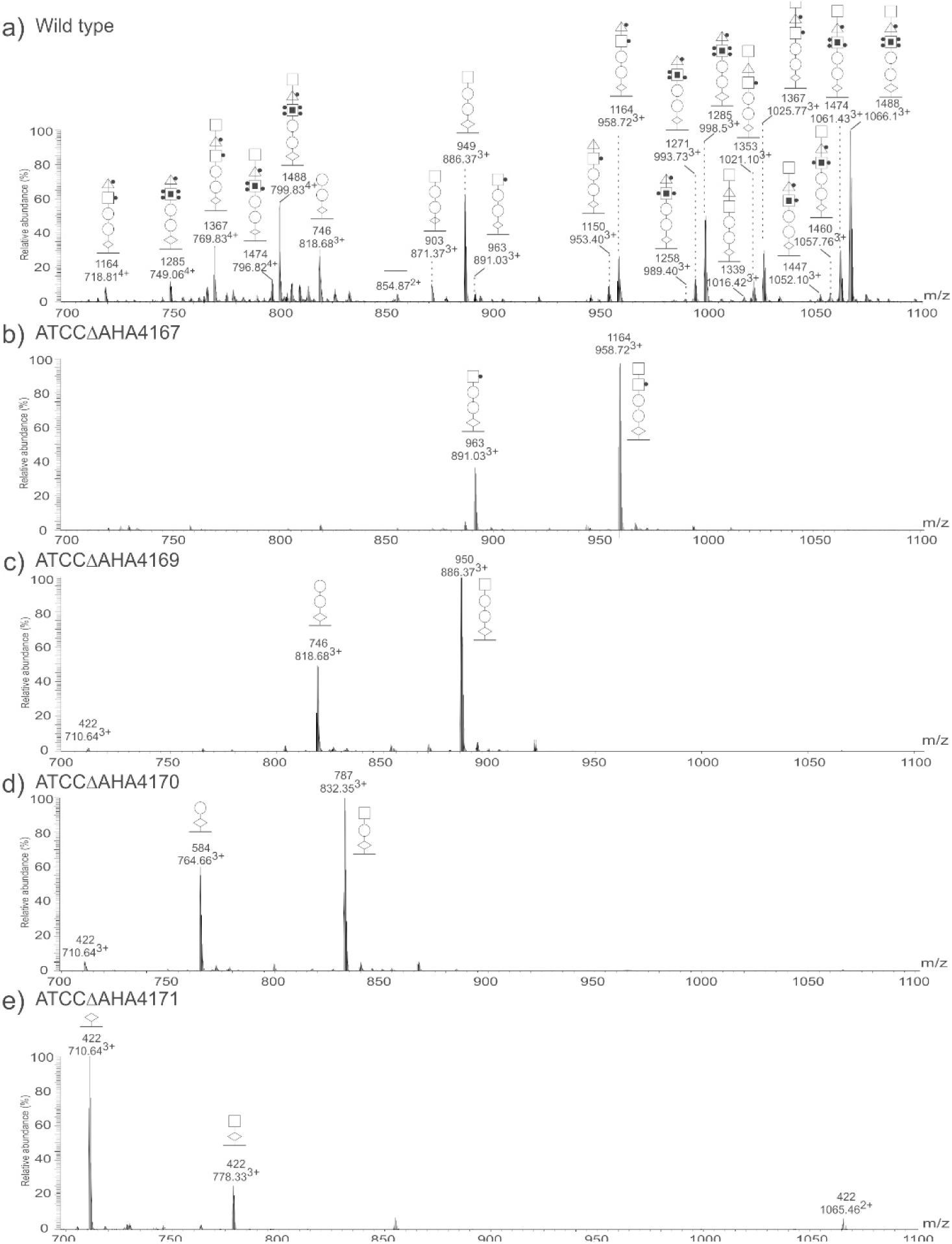
Highlighting glycan variability associated with the isobaric FlaA/FlaB T^94-108^ peptides with nLC-MS. Shown is the combined nLC-MS spectrum between 7.5 and 8.5 min, with peaks representing the 1707.6 Da FlaA/FlaB T^94-108^ isobaric peptides and associated glycosylation. Bacteria from wild type (a) modified these peptides with a highly heterologous set of penta- and hexa-saccharide glycans. Bacteria from mutant strains ATCCΔAHA4167 (b), ATCCΔAHA4169 (c), ATCCΔAHA4170 (d), and ATCCΔAHA4171 (e) do not extend the glycan beyond positions four, three, two, and one, respectively. In each mutant, however, the truncated glycan is also observed with an additional HexNAc residue. The additional HexNAc capping sugar was only observed in association with the isobaric peptides. Glycan masses (Da) are indicated above the glycopeptide ion m/z. 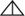: dHexNAc, 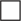: HexNAc, 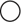: Hex, 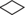: 422 Da pseudaminic acid derivative, 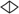: 376 Da pseudaminic acid derivative, 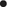:Methyl, 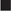: Phosphate, 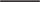: Peptide

### 3.4 Evaluation of AHA_4171 cross-complementation

The *AHA_4171* encoded protein of *A. hydrophila* ATCC 7966^T^ was a conserved glycosytransferase that showed 70% similarity and 55% identity to the Fgi-1 of *A. piscicola* AH-3 (both encoded within the FGI), although they belong to different Fgi-1 groups [16]. As described above, deletion of *AHA_4171* resulted in a truncated glycan that was not extended beyond the 422 Da Pse_d_ monosaccharide for all unique FlaA or FlaB peptides (or the disaccharide observed in association with the isobaric FlaA/FlaB peptide). We had previously demonstrated that ATCCΔAHA4171 mutant was fully complemented with its own glycosyltransferase, cloned in the pBAD33-AHA4171 plasmid [16]. However, cross-complementation of this mutant with the *A. piscicola* AH-3 *fgi-1* gene (pBAD33-Fgi-1) did not restore the ATCC7966^T^ polar flagellin molecular weight in SDS-PAGE [16]. In order to analyze if Fgi-1 of *A. piscicola* AH-3 is able to transfer any carbohydrate to the Pse_d,_ we performed LC-MS/MS analysis. The ion at m/z 828.8^2+^, selected as a representative glycopeptide, showed the 1233.62 Da FlaB T^159-166^ peptide modified with only the 422 Da Pse_d_ (the longest observed glycan) confirming that the cross-complementation did not restore the wild type glycan chain (**Fig. 7**).

**Figure 7:**
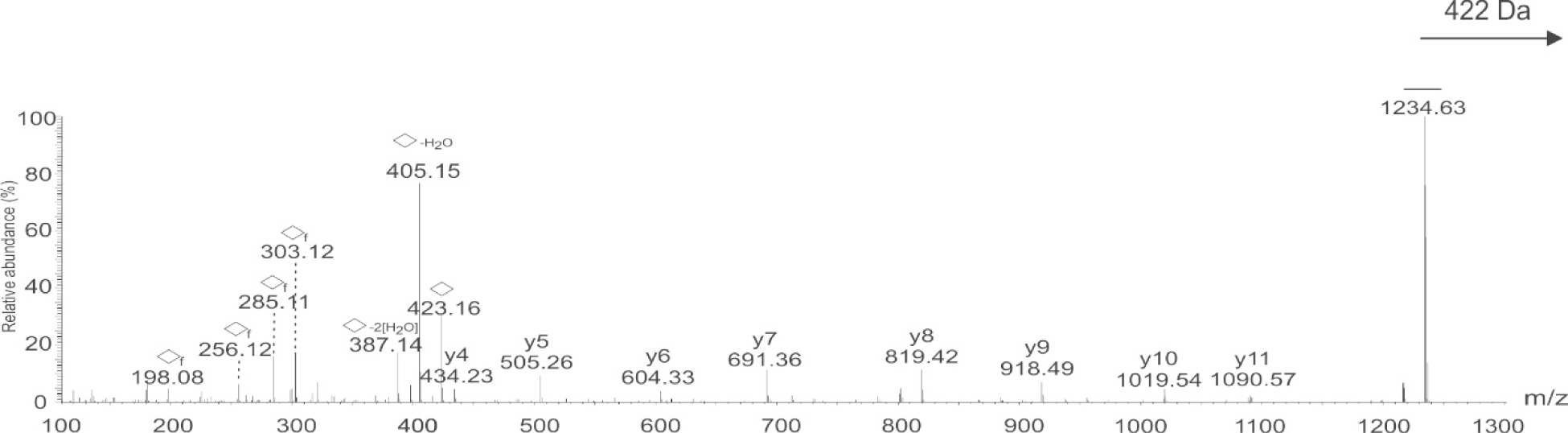
Cross-complementation of *A. hydrophila* ATCC 7966^T^ ΔAHA_4171 with *A. piscicola* AH-3 *fgi-1*. LC-MS/MS analysis of ATCCΔAHA4171 k AH-3 *fgi-1* (the 422 Da Pse derivative) revealed spectra with the same glycosylation pattern as ATCC 7966^T^ ΔAHA_4171. The cross-complementation did not restore the full glycan chain lengths. The ion at m/z 828.8^2+^ represents the 1233.62 Da FlaB T^159-166^ peptide modified with a monosaccharide as its longest glycan. Shown is the LC-MS/MS fragmentation spectrum confirming the peptide sequence and glycan composition. 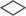: 422 Da Pse derivative, 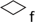: 422 Da Pse fragment, 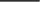: Peptide.

## 4. DISCUSSION

The glycosylation of *A. hydrophila* ATCC 7966^T^ polar flagellin structural proteins FlaA and FlaB is extremely complex, with both micro- and macro-heterogeneity previously described [18]. The glycans observed were predominantly penta- and hexa-saccharides with the basic pentasaccharide sequence being Pse_d_-Hex-Hex-MeGlcNAc-dGlcNAc and the hexasaccharide also having an additional HexNAc capping sugar. These glycans also had additional and highly variable secondary methylation and phosphorylation of the GlcNAc in the fourth position in the glycan. Most Aeromonads encode for Pse biosynthetic enzyme homologues within their genomes. The region between these genes is the FGI, which in Group II encodes for a number of putative glycosyl-, methyl, phospho-, and acetyl-transferases, as well as several lipid biosynthesis genes [16]. To better understand the polar flagellin glycan assembly in *A. hydrophila* ATCC 7966^T^, a panel of four deletion mutants were created, each lacking a putative glycosyltransferase gene located within the FGI. The flagellin glycans observed in the ATCCΔAHA4171, ATCCΔAHA4170, ATCCΔAHA4169, and ATCCΔAHA4167 mutant strains were truncated predominantly to mono-, di-, tri-, and tetra-saccharides, respectively, as compared to the penta- and hexa-saccharides observed in the wild type strain. It is worth noting that the donor and acceptor specificity of these putative glycosyltransferase enzymes appeared to be high for all four of these enzymes.

Interestingly, there was a second glycoform observed only in association with the isobaric FlaA/FlaB that had an additional HexNAc residue added to any chain length present in all of the mutant strains, creating glycan chains that were consistently one monosaccharide longer. This suggested the existence of another glycosyltransferase that has not yet been identified. This unidentified glycosyltransferase enzyme would have seemingly high donor specificity (observed to consistently transfer a HexNAc as a capping sugar onto the glycan in both wild type and mutant bacteria) but seemingly low acceptor specificity (observed to transfer the HexNAc onto any available glycan chain length regardless of the penultimate carbohydrate). The FGI of *A. hydrophila* ATCC 7966^T^ encodes several other genes, including five hypothetical genes with no known homology. It is possible that one of these genes annotated as hypothetical is in fact a novel glycosyltransferase involved in the glycan assembly, but this remains to be evaluated. Interestingly, these isobaric peptide sequences that are uniquely observed to be modified by longer glycans regardless of strain are located immediately adjacent to a toll-like receptor 5 (TLR5) binding site [32,33] in both flagellin proteins. It has been shown in *C. jejuni* that changes to the TLR5 binding site amino acid sequence compromised the stability of the flagellar filament [34], suggesting that despite its recognition by host TLR5, this sequence must be maintained in order to preserve motility and other functions of the flagellum. The modification site-specific extended glycan capped by an additional HexNAc residue observed in this study may serves to shield this pathogen from host recognition, perhaps through the steric or structural hinderance [35] afforded by the longer glycan or through host mimicry [36] since HexNAc is a carbohydrate common to both bacteria and eukaryotes. There is precedent for this in other bacteria. For example, studies of *Vibrio vulnificus* showed that its flagellin B could not stimulate TLR5 until it was deglycosylated either enzymatically or through mutagenesis of glycosylation sites [37]. Another example involved *Burkholderia cenocepacia.* It has been show that glycosylation of the flagellin in this bacterium impairs TLR5 stimulation in epithelial cells, which in turn reduced the inflammatory response [38].

The extensive postglycosylational modification of the GlcNAc occupying the fourth position of the polar flagellin glycan in wild type bacteria was not observed in any deletion mutant evaluated. Instead, this GlcNAc was consistently observed modified with a single Me group on glycans long enough to include the fourth carbohydrate. Though a single Me group was observed, it may be speculated that most of the secondary modifications are added only after the complete penta- (or hexa-) saccharide is assembled, even though phosphorylation of carbohydrates post-glycan biosynthesis is considered quite rare [39]. Further investigation into the activity of the putative methyl- and phospho-transferase gene products encoded with the FGI is therefore warranted to better understand the regulation of the secondary modification of carbohydrates.

The putative glycosyltransferase mutants ATCCΔAHA4167, ATCCΔAHA4169, ATCCΔAHA4170, and ATCCΔAHA4171 (*fgi-1*) produced polar flagellin glycans that were predominantly tetra-, tri-, di-, and mono-saccharides, respectively (excluding the exceptional isobaric peptides). The proposed involvement of each gene product is captured in **Fig. 8**. In a previous study, elimination of polar flagellin glycosylation through the deletion of *pse* biosynthesis homologues resulted in non-motile bacteria [18]. In this current study, all four glycosylation mutants, producing truncated glycans, had similarly reduced but not abolished motility compared to wild type bacteria. Therefore, motility is only possible if the flagellin proteins are at least modified by the Pse_d_ linking sugar, and full motility is only possible with a complete glycan chain.

**Figure 8:**
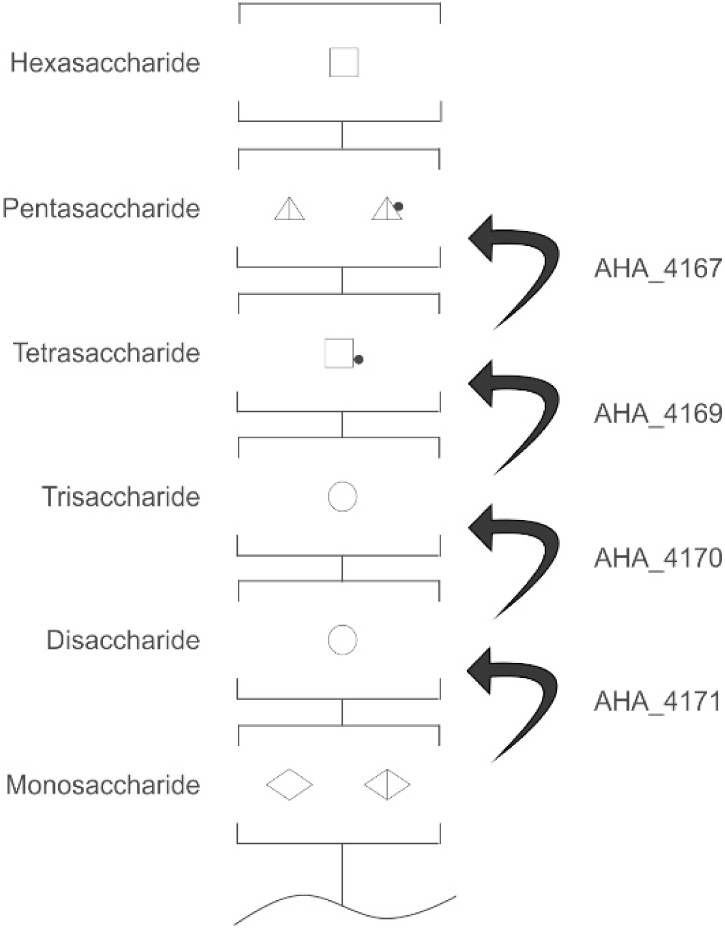
Proposed involvement of each gene in wild type polar flagellin glycan biosynthesis. 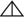: dHexNAc, 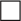: HexNAc, 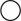: Hex, 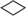: 422 Da pseudaminic acid derivative, 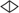: 376 Da pseudaminic acid derivative, 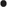:Methyl, 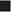: Phosphate, 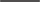: Peptide

Intermediate glycan chain lengths did not seem to impact the extent of motility that was possible. In fact, there are several species of motile bacteria, such as *Aeromonas hydrophila* AH-1 [40], that exclusively modify their flagellin protein with a single Pse_d_ monosaccharide and do not produce or require the more complex polysaccharide glycans like those observed in *A. hydrophila* ATCC 7966^T^ and *A. piscicola* AH-3. The role of Pse in modification goes beyond motility, as it has also been linked to biofilm formation, adhesion to host cells, and colonization [40–44].

In other bacteria, chaperones such as FliS in *C. jejuni* are responsible for the export of flagellin, which enables flagellum biogenesis [45]. Deletion of the gene encoding for FliS resulted in truncated or incomplete flagellar filament formation and reduced motility. It is known that flagellin glycosylation can influence the flagellin-chaperone interactions [46]. In fact, in *A. caviae,* the flagellin-specific chaperone FlaJ was shown to bind with higher affinity to glycosylated FlaA than the unmodified form of the protein. This led to more effective export of glycosylated flagellin [15] and intracellular accumulation of unglycosylated flagellin in this bacterium [15]. Intracellular accumulation of unmodified flagellin has also been observed in other species such as as *C. jejuni* [9,47] and *Bacillus subtilis* [48]. *A. caviae* modifies its flagellins with a single Pse moiety and is fully motile. Since flagellins purified from ATCCΔAHA_4171 predominantly harbour only a single Pse_d_, are still able to assemble a flagellar filament, and are at least partially motile, it is possible that Pse_d_ is the most critical feature for chaperone recognition and subsequent export of the flagellins. The variable glycan chain lengths observed across the mutant strains all had similar levels of motility, as long as the Pse_d_ monosaccharide was present, which might suggest that the longer glycans are not involved in chaperone binding, but may instead simultaneously serve other purposes, such as folding of flagellins or host immune evasion as described above. Motility is a bacterial virulence factor, with flagellin glycosylation increasingly recognized as a critical prerequisite of the virulence factors. Preferential chaperone-mediated export of glycosylated flagellin conceptually links this posttranslational modification to the final structure and function.

The FGI of *A. piscicola* AH-3 contains the *fgi-1* glycosyltransferase gene, with activity of the gene product confirmed previously [16]. Deletion of the *fgi-1* gene resulted in polar flagellin glycans that were truncated to a single monosaccharide modification and a significant reduction in motility. The FGI of *A. hydrophila* ATCC 7966^T^ encodes a homologue of this gene, denoted *AHA_4171*. In the previous study, despite the homology, cross-complementation of *A. piscicola* AH3Δ*fgi-1* with *AHA_4171* did not restore glycan assembly or motility to wild type levels.

Similarly, in this study, cross-complementation of ATCCΔAHA4171 with the *fgi-1* gene from *A. piscicola* AH-3 did not restore glycan assembly or motility. Both enzymes appear to transfer a Hex onto the Pse derivative in their respective species. The two Pse derivatives are related but different structures, with that of AH-3 being 376 Da and that of ATCC 7966^T^ being 422 Da. Furthermore, the stereochemistry of the hexose residues in both species have not been assigned due to a lack of observable oxonium ion that would permit MS^3^ fragmentation and further structural elucidation. Therefore, the inability to cross-complement may result from donor or acceptor specificity of the glycosyltransferases, or even perhaps both. Beyond improving understanding of this infectious disease, the identification of novel glycosyltransferase enzymes with high substrate specificity is useful for expanding the tools available for glycoengineering. There are many applications of glycoengineering [49], including improving protein-based therapeutics by increasing efficacy, reducing side-effects, or extending half-life [50].

Since disruption of flagellin glycosylation leads to reduced motility in *A. hydrophila* ATCC^T^ 7966, as in many other bacteria, improved understanding of flagellin glycan biosynthesis and structures may unveil avenues for novel therapeutic development [51,52]. Though only evaluated *in vitro* to date, a few potential antivirulence strategies have been explored. Small molecule inhibitors of the *C. jejuni* and *Helicobacter pylori* glycan biosynthetic pathway have been shown to block flagellar filament production *in vitro*, for example [53]. The motility of *C. jejuni* was also successfully reduced by the introduction of *Campylobacter* bacteriophage NCTC 12673 protein FlaGrab, which is known to specifically bind to the Pse5Ac7Am sugar modifying the *C. jejuni* flagellin [54]. Continued investigation into the FGI of *A. hydrophila* ATCC 7966^T^, as well as flagellin glycan biosynthesis in other important human pathogens, is warranted in the pursuit of novel treatment options as antimicrobial resistance continues to spread.

## 5. CONCLUSIONS

*A. hydrophila* ATCC 7966^T^ has extensive polar flagellin glycan metaheterogeneity with complex polysaccharide glycans. This study sought to elucidate the biosynthetic pathway responsible by evaluating four gene products with homology to known glycosyltransferases located within the *A. hydrophila* ATCC 7966^T^ flagellar glycosylation island (FGI). Deletion of genes *AHA_4167*, *AHA_4179*, *AHA_4170*, and *AHA_4171* were observed to have truncated polar flagellin glycans with sequentially shorter chain lengths, respectively, and are therefore confirmed to have involvement in glycan biosynthesis. All of these mutant strains had reduced motility compared to wild type bacteria, further supporting the role of flagellin glycosylation as a perquisite for flagellar-mediated motility.

## 6. DATA AVAILABILITY STATEMENT

The publicly available NCBI RefSeq assembly for *A. hydrophila* subsp. *hydrophila* ATCC 7966^T^ (GCF_000014805.1) was used for protein identification. Mass spectrometry files generated will be made available upon reasonable request by contacting the corresponding author.

## 7. ACKNOWLEDGEMENTS

This work was partially supported by the Spanish Ministerio de Ciencia e Innovación, Plan Nacional de I + D with grant number PID2021-124676NB-I00. The authors gratefully acknowledge the funding of the Pilot Mass Spectrometry Collaboration Center by Laboratories Canada. J.C.S. acknowledges the support of the Natural Sciences and Engineering Research Council of Canada. The authors wish to thank Maite Polo for her technical assistance.

